# A highly conserved cryptic epitope in the receptor-binding domains of SARS-CoV-2 and SARS-CoV

**DOI:** 10.1101/2020.03.13.991570

**Authors:** Meng Yuan, Nicholas C. Wu, Xueyong Zhu, Chang-Chun D. Lee, Ray T. Y. So, Huibin Lv, Chris K. P. Mok, Ian A. Wilson

## Abstract

The outbreak of COVID-19, which is caused by SARS-CoV-2 virus, continues to spread globally, but there is currently very little understanding of the epitopes on the virus. In this study, we have determined the crystal structure of the receptor-binding domain (RBD) of the SARS-CoV-2 spike (S) protein in complex with CR3022, a neutralizing antibody previously isolated from a convalescent SARS patient. CR3022 targets a highly conserved epitope that enables cross-reactive binding between SARS-CoV-2 and SARS-CoV. Structural modeling further demonstrates that the binding site can only be accessed when at least two RBDs on the trimeric S protein are in the “up” conformation. Overall, this study provides structural and molecular insight into the antigenicity of SARS-CoV-2.

**ONE SENTENCE SUMMARY:** Structural study of a cross-reactive SARS antibody reveals a conserved epitope on the SARS-CoV-2 receptor-binding domain.

## MAIN

The ongoing outbreak of Coronavirus Disease 2019 (COVID-19) originally emerged in China during December 2019 (*1*) and has now spread over 120 countries as of March 12, 2020 and become pandemic. COVID-19 is caused by a novel coronavirus, severe acute respiratory syndrome coronavirus 2 (SARS-CoV-2) (*2*). In fact, two other coronaviruses have caused global outbreak in the past two decades, namely SARS-CoV (2002-2003) and Middle East respiratory syndrome coronavirus (MERS-CoV) (2012-present). The surface spike glycoprotein (S), which is critical for virus entry through engaging the host receptor and mediating virus-host membrane fusion, is the major antigen of coronaviruses. The S proteins of SARS-CoV-2 and SARS-CoV, which are phylogenetically closely related, have an amino-acid sequence identity of around 77% (*3*). Such a high degree of sequence similarity raises the possibility that cross-reactive epitopes may exist. A recent study has shown that CR3022, which is a human neutralizing antibody that targets the receptor-binding domain (RBD) of SARS-CoV (*4*), can bind to the RBD of SARS-CoV-2 (*5*). This finding provides an opportunity to uncover a cross-reactive epitope.

CR3022 was previously isolated from a convalescent SARS patient and is encoded by germline genes IGHV5-51, IGHD3-10, IGHJ6 (heavy chain), and IGKV4-1, IGKJ2 (light chain) (*4*). Based on IgBlast analysis (*6*), the IGHV of CR3022 is 3.1% somatically mutated at the nucleotide sequence level, which results in eight amino-acid changes from the germline sequence, whereas IGKV of CR3022 is 1.3% somatically mutated resulting in three amino-acid changes from the germline sequence (fig. S1). We therefore determined the crystal structure of CR3022 with the SARS-CoV-2 RBD at 3.1 Å resolution (table S1 and fig.S2, A and B) (*7*). CR3022 uses both heavy and light chains (Fig. 1B), and all six complementarity-determining region (CDR) loops (Fig. 1C) for interaction with the RBD. The buried surface area on the epitope is 917 Å^2^ and SARS-CoV-2 recognition by CR3022 is largely driven by hydrophobic interactions (Fig. 1E). Five out of 11 somatic mutations are found in the paratope region (fig. S2C), implying their likely importance in the affinity maturation process.

**Fig. 1.**
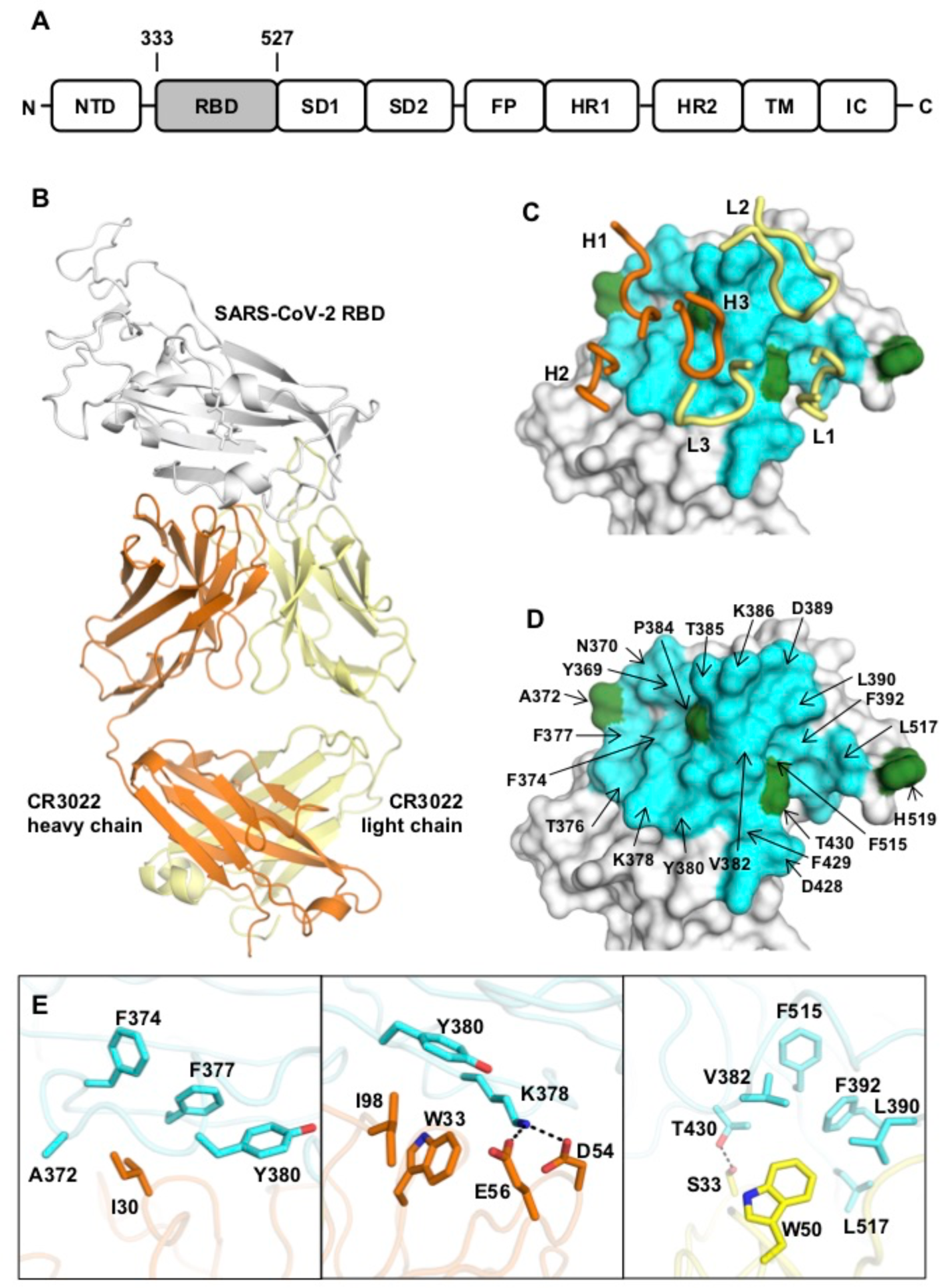
Crystal structure of CR3022 in complex with SARS-CoV-2 RBD. **(A)** Overall topology of the SARS-CoV-2 spike glycoprotein. NTD: N-terminal domain, RBD: receptor-binding domain, SD1: subdomain 1, SD2: subdomain 2, FP: fusion peptide, HR1: heptad repeat 1, HR2: heptad repeat 2, TM: transmembrane region, IC: intracellular domain. **(B)** Structure of CR3022 Fab in complex with SARS-CoV-2 RBD. CR3022 heavy chain is colored in orange, CR3022 light chain in yellow, and SARS-CoV-2 RBD in light grey. **(C-D)** Epitope residues on SARS-CoV-2 are colored in cyan and green. CDR loops are labeled. Cyan: epitope residues that are conserved between SARS-CoV-2 and SARS-CoV. Green: epitope residues that are not conserved between SARS-CoV-2 and SARS-CoV. **(D)** Epitope residues that are important for binding to CR3022 are labeled. Epitope residues are defined here as residues in SARS-CoV-2 RBD with buried surface area > 0 Å^2^ after Fab CR3022 binding as calculated with PISA (*35*). **(E)** Several key interactions between CR3022 and SARS-CoV-2 RBD are highlighted. CR3022 heavy chain is colored in orange, CR3022 light chain in yellow, and SARS-CoV-2 RBD in cyan. Hydrogen bonds are represented by dashed lines.

Out of 28 residues in the epitope (defined as residues buried by CR3022), 24 (86%) are conserved between SARS-CoV-2 and SARS-CoV (Fig. 1D). This high sequence conservation explains the cross-reactivity of CR3022. Nonetheless, despite having a high conservation in the epitope residues, CR3022 Fab binds to SARS-CoV RBD (K_d_ = 1 nM) with a much higher affinity than to SARS-CoV-2 RBD (K_d_ = 115 nM) (Table 1 and fig. S3). We postulate that the difference in binding affinity of CR3022 between SARS-CoV-2 and SARS-CoV RBDs is due to the non-conserved residues in the epitope (fig. S4). The most dramatic difference between the CR3022 epitope in SARS-CoV-2 and SARS-CoV is an additional N-glycosylation site at N370 (N357 in SARS-CoV numbering). The N-glycan sequon (NxS/T) arises from an amino-acid difference at residue 372, where SARS-CoV has a Thr compared to Ala in SARS-CoV-2 (fig. S4B). Mass spectrometry analysis has shown that a complex glycan is indeed present at this N-glycosylation site in SARS-CoV (*8*). An N-glycan at N370 would fit into a groove formed between heavy and light chains (fig. S4C), which could increase contact and, hence, binding affinity to CR3022. We then tested whether CR3022 was able to neutralize SARS-CoV-2 and SARS-CoV in an *in vitro* microneutralization assay (*7*). While CR3022 could neutralize SARS-CoV, it did not neutralize SARS-CoV-2 at the highest concentration tested (400 µg/mL) (fig. S5). This *in vitro* neutralization result is consistent with lower affinity binding of CR3022 for SARS-CoV-2, although other explanations are possible as outlined below.

**Table 1.** Binding affinity of CR3022 to recombinant RBD and S protein.

| **Affinity ($K_d$ in nM)** | **CR3022 IgG** | **CR3022 Fab** |
| --- | --- | --- |
| SARS-CoV-2 RBD | < 0.1 | 115 ± 3 |
| SARS-CoV RBD | < 0.1 | 1.0 ± 0.1 |

SARS-CoV-2 uses the same host receptor, angiotensin I converting enzyme 2 (ACE2) as SARS-CoV (*3, 9-11*). Interestingly, the epitope of CR3022 does not overlap with the ACE2-binding site (Fig. 2A). Structural alignment of CR3022-SARS-CoV-2 RBD complex with the ACE2-SARS-CoV-2 RBD complex (*11*) further indicates that binding of CR3022 would not clash with ACE2 (*12*). This analysis implies that the neutralization mechanism of CR3022 does not depend on direct blocking of receptor binding, which is consistent with the observation that CR3022 does not compete with ACE2 for binding to the RBD (*5*). Unlike CR3022, most known SARS RBD-targeted antibodies compete with ACE2 for binding to RBD (*4, 13-16*). The epitopes of these antibodies are very different from that of CR3022 (Fig. 2B). In fact, it has been shown that CR3022 can synergize with other RBD-targeted antibodies to neutralize SARS-CoV (*4*). Although CR3022 itself cannot neutralize SARS-CoV-2 in this *in vitro* assay, whether CR3022 can synergize with other SARS-CoV-2 RBD-targeted monoclonal antibodies for neutralization remains to be determined.

**Fig. 2.**
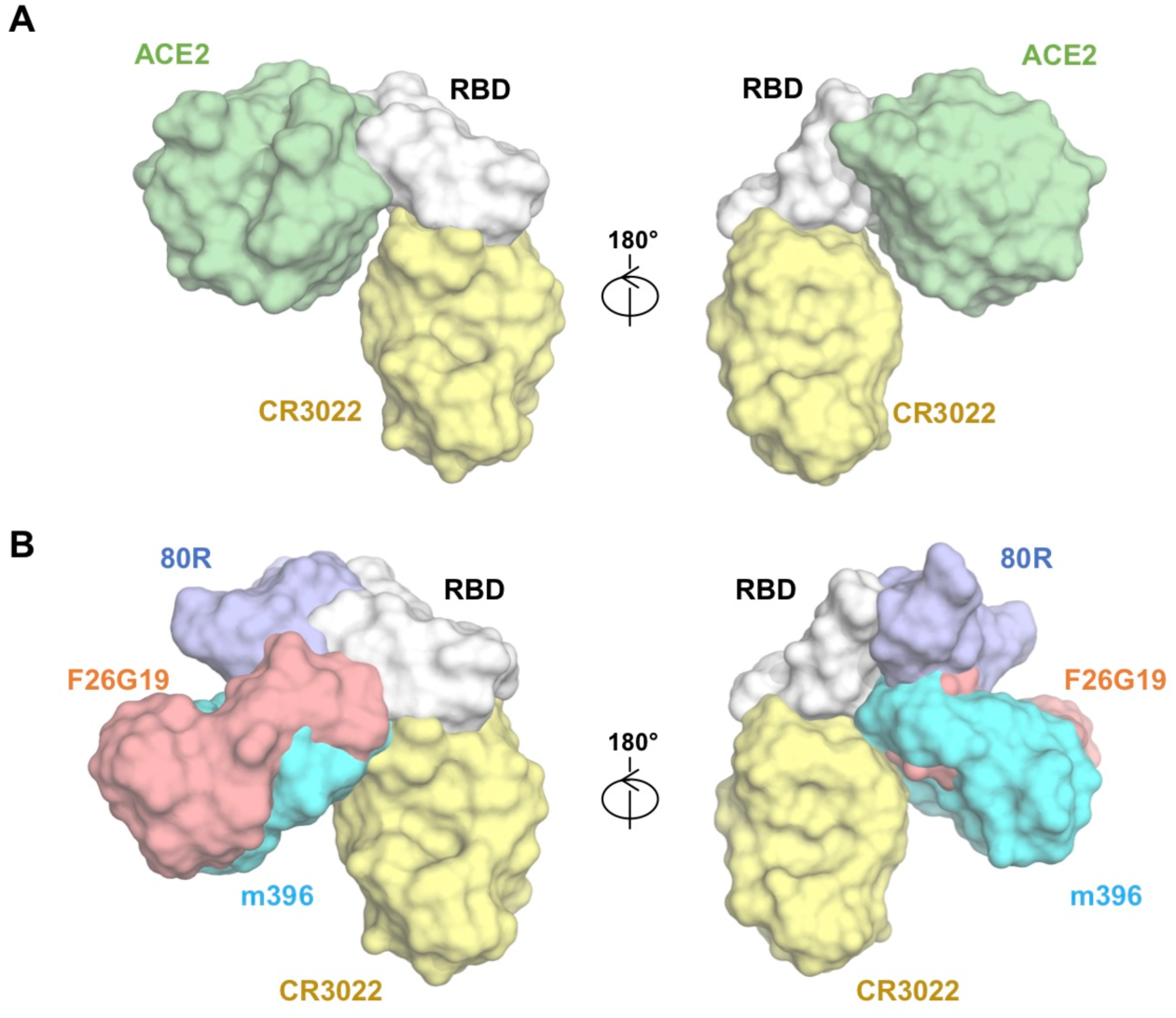
The relative binding position of CR3022 with respect to receptor ACE2 and other SARS-CoV RBD monoclonal antibodies. **(A)** Structures of CR3022-SARS-CoV-2 RBD complex and ACE2-SARS-CoV-2 RBD complex (*11*) are aligned based on the SARS-CoV-2 RBD. ACE2 is colored in green, RBD in light grey, and CR3022 in yellow. **(B)** Structural superposition of CR3022-SARS-CoV-2 RBD complex, F26G19-SARS-CoV RBD complex (PDB 3BGF) (*36*), 80R-SARS-CoV RBD complex (PDB 2GHW) (*37*), and m396-SARS-CoV RBD complex (PDB 2DD8) (*16*).

Recently, the cryo-EM structure of homotrimeric SARS-CoV-2 S protein was determined (*17, 18*) and demonstrated that the RBD, as in other coronaviruses (*19, 20*) adopts two different dispositions in the trimer. The RBD can then undergo a hinge-like movement to transition between “up” or “down” conformations (Fig. 3A). ACE2 host receptor can only interact with the RBD when it is in the “up” conformation, whereas the “down” conformation is inaccessible to ACE2. Interestingly, the epitope of CR3022 is also only accessible when the RBD is in the “up” conformation (Fig. 3, B and C). Furthermore, the ability for CR3022 to access the RBD also depends on the relative disposition of the RBD on the adjacent protomer. CR3022 can only access RBD when the targeted RBD on one protomer of the trimer and the RBD on the adjacent protomer are both in the “up” conformation. The variable region of CR3022 would clash with the RBD on the adjacent protomer if the latter adopts a “down” conformation (Fig. 3D). As a homotrimer, the S protein could potentially adopt four possible RBD configurations, namely none-”up”, single-”up”, double-”up”, and triple-”up”. It appears that CR3022 can only bind to the S protein when it is in double-”up” or triple-”up” configuration. Specifically, one molecule of CR3022 can be accommodated in the double-”up” configuration (Fig. 3E), whereas three molecules of CR3022 could potentially be accommodated in the triple-”up” configuration (Fig. 3F). Previous cryo-EM studies have also shown that the recombinant SARS-CoV S protein is mostly found in the none-”up”, single-”up”, or double-”up” conformations (*19, 21*), but rarely in the triple-”up” conformation, even with ACE2 receptor bound (*21, 22*). Together with the fact that CR3022 was isolated from a convalescent SARS patient (*4*), these observations suggest that an antibody response can be elicited against this cryptic epitope in SARS-CoV and possibly SARS-CoV-2. Structural comparison shows that, as compared to SARS-CoV-2, the “up” conformation of RBD in SARS-CoV has a larger dihedral angle to the horizontal plane of the S protein (fig. S6), suggesting that the CR3022 epitope may be slightly more accessible in SARS-CoV than in SARS-CoV-2, in these spike ectodomain constructs. Nevertheless, the availability of this cryptic epitope on the actual virus surface still has to be quantified to fully comprehend its immunological role during natural infection.

**Fig. 3.**
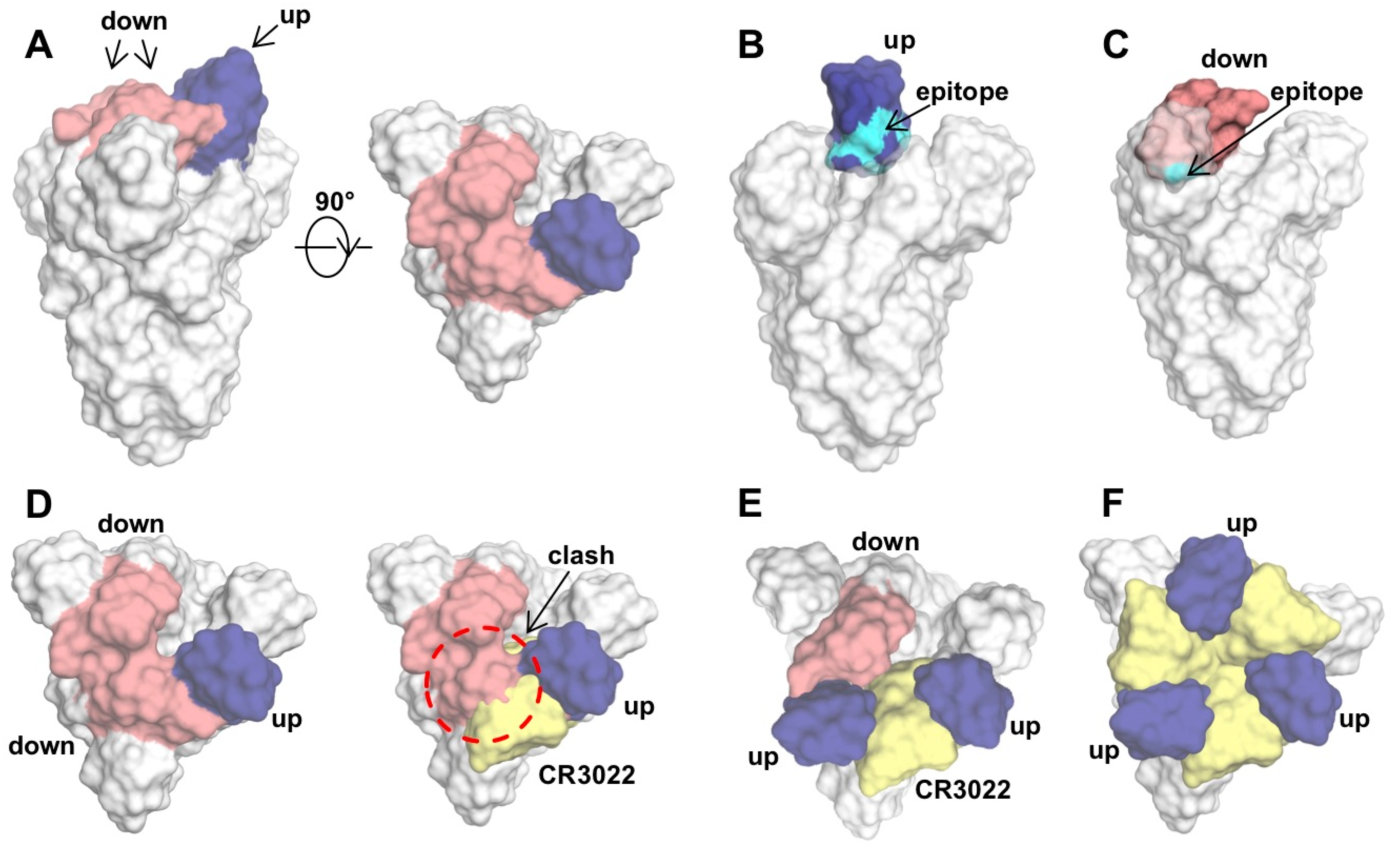
Binding of CR3022 depends on the RBD configurations on the S protein. **(A)** RBD in the S proteins of SARS-CoV-2 and SARS-CoV can adopt either an “up” conformation (blue) or a “down” conformation (red). PDB 6VSB (cryo-EM structure of SARS-CoV-2 S protein) (*17*) is shown. **(B-C)** CR3022 epitope (cyan) on the RBD is exposed in **(B)** the “up” but not **(C)** the “down” conformation. **(D)** Binding of CR3022 to single-”up” configuration would clash (indicated by the red circle) with the neighboring RBD. CR3022 is colored yellow. **(E-F)** The double-”up” and triple-”up” RBD configurations on the SARS-CoV-2 S protein are modeled based on PDB 6VSB (*17*). **(E)** One CR3022 molecule can be accommodated per S protein in the double-”up” configuration, and **(F)** three CR3022 molecules could potentially be accommodated per S protein in the triple-”up” configuration.

Overall, our study provides insight into how SARS-CoV-2 can be targeted by the humoral immune response and revealed a conserved, but cryptic epitope shared between SARS-CoV-2 and SARS-CoV. Recently, our group and others have identified a conserved epitope on influenza A virus hemagglutinin (HA) that is located in the trimeric interface and is only exposed through protein “breathing” (*23-25*), which is somewhat analogous to the epitope of CR3022. Antibodies to this influenza HA trimeric interface epitope do not exhibit *in vitro* neutralization activity but can confer *in vivo* protection. Similarly, antibodies to another conserved epitope that partially overlaps with the influenza HA trimeric interface also are non-neutralizing *in vitro*, but protective *in vivo* (*26*). Furthermore, there are other examples of antibodies that do not have *in vitro* neutralization activity but confer *in vivo* protection, such as reported for influenza virus (*27*), herpesvirus (*28*), cytomegalovirus (*29*), alphavirus (*30*), and dengue virus (*31*). Therefore, although CR3022 does not neutralize SARS-CoV-2 *in vitro* despite its reasonable binding affinity, it is possible that this epitope can confer *in vivo* protection. The potential existence of non-neutralizing protective antibodies to SARS-CoV-2 highlights the need for an effective SARS-CoV-2 infection mouse model, which has yet to be established.

Since there is currently great urgency in the efforts to develop a vaccine against SARS-CoV-2, characterizing the epitopes on SARS-CoV-2 S protein is extremely valuable. Much work is now ongoing in isolating human monoclonal antibodies from SARS-CoV-2 patients. We anticipate that these investigations will decipher the antigenic properties and major epitopes of SARS-CoV-2. In addition, the molecular features that antibodies use for targeting neutralizing epitopes can inspire development of therapeutics, such as antiviral peptides and small molecules (*32*). As this coronavirus outbreak continues to pose an enormous global risk (*33, 34*), the availability of conserved epitopes may allow structure-based design not only of a SARS-CoV-2 vaccine, but also for cross-protective antibody responses against future coronavirus epidemics and pandemics. While a more universal coronavirus vaccine is not the most urgent goal at present, it is certainly worthwhile for future consideration especially as cross-protective epitopes are identified so that we can be better prepared for the next novel coronavirus outbreak.

## ACKNOWLEDGEMENTS

We thank Henry Tien for technical support with the crystallization robot, Jeanne Matteson for contribution to mammalian cell culture, Wenli Yu to insect cell culture, Robyn Stanfield for assistance in data collection, and Andrew Ward for discussion.

## Author contributions

M.Y., N.C.W., X.Z. and I.A.W. conceived and designed the study. M.Y., N.C.W. and C.C.D.L. expressed and purified the proteins. M.Y. and N.C.W. performed biolayer interferometry binding assays. R.T.Y.S., H.L. and C.K.P.M. performed the neutralization experiments. M.Y., N.C.W. and X.Z. collected the X-ray data, determined and refined the X-ray structures. M.Y., N.C.W. and C.K.P.M. analyzed the data. M.Y., N.C.W. and I.A.W. wrote the paper and all authors reviewed and edited the paper.

## Data and materials availability

X-ray coordinates and structure factors are deposited at the RCSB Protein Data Bank under accession code: 6W41. All of the other data that support the conclusions of the study are available from the corresponding author upon request.

## REFERENCES AND NOTES

1. C. Huang et al., Clinical features of patients infected with 2019 novel coronavirus in Wuhan, China. Lancet 395, 497–506 (2020).

2. V. Coronaviridae Study Group of the International Committee on Taxonomy of, The species Severe acute respiratory syndrome-related coronavirus: classifying 2019-nCoV and naming it SARS-CoV-2. Nat Microbiol 10.1038/s41564-020-0695-z (2020).

3. P. Zhou et al., A pneumonia outbreak associated with a new coronavirus of probable bat origin. Nature 10.1038/s41586-020-2012-7 (2020).

4. J. ter Meulen et al., Human monoclonal antibody combination against SARS coronavirus: synergy and coverage of escape mutants. PLoS Med 3, e237 (2006).

5. X. Tian et al., Potent binding of 2019 novel coronavirus spike protein by a SARS coronavirus-specific human monoclonal antibody. Emerg Microbes Infect 9, 382–385 (2020).

6. J. Ye, N. Ma, T. L. Madden, J. M. Ostell, IgBLAST: an immunoglobulin variable domain sequence analysis tool. Nucleic Acids Res 41, W34–40 (2013).

7. See supplementary materials.

8. Y. Watanabe et al., Vulnerabilities in coronavirus glycan shields despite extensive glycosylation. bioRxiv 10.1101/2020.02.20.957472 (2020).

9. M. Letko, A. Marzi, V. Munster, Functional assessment of cell entry and receptor usage for SARS-CoV-2 and other lineage B betacoronaviruses. Nat Microbiol 10.1038/s41564-020-0688-y (2020).

10. R. Yan et al., Structural basis for the recognition of the SARS-CoV-2 by full-length human ACE2. Science 10.1126/science.abb2762 (2020).

11. J. Lan et al., Crystal structure of the 2019-nCoV spike receptor-binding domain bound with the ACE2 receptor. bioRxiv 10.1101/2020.02.19.956235 (2020).

12. ACE2 also forms a dimer when it associates with an amino acid transporter B^0^AT1 (10). We modeled a CR3022 IgG onto this dimer structure and found no clashes of CR3022 with ACE2 in its dimeric form where the RBDs would likely come from adjacent trimers on the virus (10).

13. J. Sui et al., Potent neutralization of severe acute respiratory syndrome (SARS) coronavirus by a human mAb to S1 protein that blocks receptor association. Proc Natl Acad Sci U S A 101, 2536–2541 (2004).

14. E. N. van den Brink et al., Molecular and biological characterization of human monoclonal antibodies binding to the spike and nucleocapsid proteins of severe acute respiratory syndrome coronavirus. J Virol 79, 1635–1644 (2005).

15. J. D. Berry et al., Development and characterisation of neutralising monoclonal antibody to the SARS-coronavirus. J Virol Methods 120, 87–96 (2004).

16. P. Prabakaran et al., Structure of severe acute respiratory syndrome coronavirus receptor-binding domain complexed with neutralizing antibody. J Biol Chem 281, 15829–15836 (2006).

17. D. Wrapp et al., Cryo-EM structure of the 2019-nCoV spike in the prefusion conformation. Science 10.1126/science.abb2507 (2020).

18. A. C. Walls et al., Structure, function, and antigenicity of the SARS-CoV-2 spike glycoprotein. Cell 10.1016/j.cell.2020.02.058 (2020).

19. Y. Yuan et al., Cryo-EM structures of MERS-CoV and SARS-CoV spike glycoproteins reveal the dynamic receptor binding domains. Nat Commun 8, 15092 (2017).

20. M. Gui et al., Cryo-electron microscopy structures of the SARS-CoV spike glycoprotein reveal a prerequisite conformational state for receptor binding. Cell Res 27, 119–129 (2017).

21. R. N. Kirchdoerfer et al., Stabilized coronavirus spikes are resistant to conformational changes induced by receptor recognition or proteolysis. Sci Rep 8, 15701 (2018).

22. Yuan et al. (19) observed 56% of the wild-type recombinant SARS-CoV S protein particle in none-”up” conformation and 44% in single-”up” conformation, while Kirchdoerfer et al. (21) found that recombinant SARS-CoV S protein, with K968P/V969P mutations in the S2 subunit to stabilize the prefusion conformation, has 58% single-”up”, 39% in double-”up”, and 3% in triple-”up” conformations. However, it is not known whether the distribution of different configurations of S proteins on virus surface is the same as that of recombinant S protein.

23. S. Bangaru et al., A site of vulnerability on the influenza virus hemagglutinin head domain trimer interface. Cell 177, 1136–1152.e1118 (2019).

24. A. Watanabe et al., Antibodies to a conserved influenza head interface epitope protect by an IgG subtype-dependent mechanism. Cell 177, 1124–1135.e1116 (2019).

25. G. Bajic et al., Influenza antigen engineering focuses immune responses to a subdominant but broadly protective viral epitope. Cell Host Microbe 25, 827–835.e826 (2019).

26. J. Lee et al., Molecular-level analysis of the serum antibody repertoire in young adults before and after seasonal influenza vaccination. Nat Med 22, 1456–1464 (2016).

27. C. Dreyfus et al., Highly conserved protective epitopes on influenza B viruses. Science 337, 1343–1348 (2012).

28. C. Petro et al., Herpes simplex type 2 virus deleted in glycoprotein D protects against vaginal, skin and neural disease. eLife 4, e06054 (2015).

29. A. Bootz et al., Protective capacity of neutralizing and non-neutralizing antibodies against glycoprotein B of cytomegalovirus. PLoS Pathog 13, e1006601 (2017).

30. C. W. Burke et al., Human-like neutralizing antibodies protect mice from aerosol exposure with western equine encephalitis virus. Viruses 10, 147 (2018).

31. E. A. Henchal, L. S. Henchal, J. J. Schlesinger, Synergistic interactions of anti-NS1 monoclonal antibodies protect passively immunized mice from lethal challenge with dengue 2 virus. J Gen Virol 69 (Pt 8), 2101–2107 (1988).

32. N. C. Wu, I. A. Wilson, Structural insights into the design of novel anti-influenza therapies. Nat Struct Mol Biol 25, 115–121 (2018).

33. V. D. Menachery et al., A SARS-like cluster of circulating bat coronaviruses shows potential for human emergence. Nat Med 21, 1508–1513 (2015).

34. V. D. Menachery et al., SARS-like WIV1-CoV poised for human emergence. Proc Natl Acad Sci U S A 113, 3048–3053 (2016).

35. E. Krissinel, K. Henrick, Inference of macromolecular assemblies from crystalline state. J Mol Biol 372, 774–797 (2007).

36. J. E. Pak et al., Structural insights into immune recognition of the severe acute respiratory syndrome coronavirus S protein receptor binding domain. J Mol Biol 388, 815–823 (2009).

37. W. C. Hwang et al., Structural basis of neutralization by a human anti-severe acute respiratory syndrome spike protein antibody, 80R. J Biol Chem 281, 34610–34616 (2006).

