## Supplementary Materials for "A highly conserved cryptic epitope in the receptor-binding domains of SARS-CoV-2 and SARS-CoV"

**This PDF file includes:**

Materials and Methods

Figs. S1 to S6

Tables S1 to S3

References 38-47

### MATERIALS AND METHODS

#### Expression and purification of RBD

The receptor-binding domain (RBD) (residues 319-541) of the SARS-CoV-2 spike (S) protein (GenBank: QHD43416.1), as well as the RBD (residues 306-527) of the SARS-CoV S protein (GenBank: ABF65836.1), were cloned into a customized pFastBac vector (*38*). The RBD constructs were fused with an N-terminal gp67 signal peptide and a C-terminal $His_6$ tag. Recombinant bacmid DNA was generated using the Bac-to-Bac system (Life Technologies). Baculovirus was generated by transfecting purified bacmid DNA into Sf9 cells using FuGENE HD (Promega), and subsequently used to infect suspension cultures of High Five cells (Life Technologies) at an MOI of 5 to 10. Infected High Five cells were incubated at 28 °C with shaking at 110 r.p.m. for 72 h for protein expression. The supernatant was then concentrated using a 10 kDa MW cutoff Centramate cassette (Pall Corporation). The S and RBD proteins were purified by Ni-NTA, followed by size exclusion chromatography, and buffer exchanged into 20 mM Tris-HCl pH 7.4 and 150 mM NaCl.

#### Expression and purification of CR3022 Fab and IgG

The CR3022 Fab heavy (GenBank: DQ168569.1) and light (GenBank: DQ168570.1) chains were cloned into phCMV3. The plasmids were transiently co-transfected into Expi293F cells at a ratio of 2:1 (HC:LC) using ExpiFectamine™ 293 Reagent (Thermo

Fisher Scientific) according to the manufacturer's instructions. The supernatant was collected at 7 days post-transfection. The Fab was purified with a CaptureSelect™ CH1-XL Pre-packed Column (Thermo Fisher Scientific) followed by size exclusion chromatography.

For full-length IgGs of CR3022, m396 (sequences from PDB 2DD8 (*16*)), and S230.15 (sequences from PDB 6NB6 (*39*)), the heavy-chain and light-chain plasmids were transiently co-transfected into ExpiCHO cells at a ratio of 2:1 using ExpiFectamine™ CHO Reagent (Thermo Fisher Scientific) according to the manufacturer's instructions. The supernatant was collected at 14 days post-transfection. The IgGs were purified using a Protein G column (GE Healthcare), and further purified by size exclusion chromatography.

**Crystallization and structural determination**

Purified CR3022 Fab and SARS-CoV-2 RBD were mixed at a molar ratio of 1:1 and incubated overnight at 4 °C. The complex (15 mg/ml) was screened for crystallization using the 384 conditions of the JCSG Core Suite (Qiagen) at 293 K on our custom-designed robotic CrystalMation system (Rigaku) at Scripps Research by the vapor diffusion method in sitting drops containing 0.1 μl of protein and 0.1 μl of reservoir solution. Optimized crystals were then grown in 80 mM sodium acetate pH 4.6, 1.5 M ammonium sulfate, and 20% glycerol. Crystals were grown for 14 days and then flash cooled in liquid nitrogen. Diffraction data were collected at cryogenic temperature (100 K) at beamline 23-ID-D of the Argonne Photon Source (APS) with a beam wavelength of 1.033 Å, and processed with HKL2000 (*40*). Structures were solved by molecular replacement using PHASER with homology models for Fab CR3022 generated from PDB ID: 4KMT (*41*) and for SARS-CoV-2 RBD generated from a structure of SARS-

CoV-RBD (PDB ID: 2AJF) (*42*) with SWISS-MODEL (*43*). Iterative model building and refinement were carried out in COOT (*44*) and PHENIX (*45*), respectively. Epitope and paratope residues, as well as their interactions, were identified by accessing PISA at the European Bioinformatics Institute (http://www.ebi.ac.uk/pdbe/prot_int/pistart.html) (*35*).

#### Biolayer interferometry binding assay

Binding assays were performed by biolayer interferometry (BLI) using an Octet Red instrument (FortéBio) as described previously (*46*). Briefly, $His_6$-tagged S and RBD proteins at 20 to 100 μg/mL in 1x kinetics buffer (1x PBS, pH 7.4, 0.01% BSA and 0.002% Tween 20) were loaded onto Anti-Penta-HIS (HIS1K) biosensors and incubated with the indicated concentrations of CR3022 Fab or IgG. The assay consisted of five steps: 1) baseline: 60 s with 1x kinetics buffer; 2) loading: 300 s with $his_6$-tagged S or RBD proteins; 3) baseline: 60 s with 1x kinetics buffer; 4) association: 120 s with samples (Fab or IgG); and 5) dissociation: 120 s with 1x kinetics buffer. For estimating the exact $K_d$, a 1:1 binding model was used.

#### Microneutralization assay

Monoclonal antibodies were mixed with equal volumes of SARS-CoV or SARS-CoV-2 at a dose of 100 tissue culture infective doses 50% ($TCID_{50}$) determined by Vero and Vero E6 cells respectively. After 1 h of incubation at 37°C, 35 μl of the virus-antibody mixture was added in quadruplicate to Vero or Vero E6 cell monolayers in 96-well microtiter plates. After 1 h of adsorption, the virus-antibody mixture was removed and replaced with 150 μl of virus growth medium in each well. The plates were incubated for 3 days at 37°C in 5% $CO_2$ in a humidified incubator. A cytopathic effect was observed at day 3 post-inoculation. The highest plasma dilution that protected 50% of the replicate wells was denoted as the neutralizing antibody titer. A virus back-titration of the input virus was included in each batch of tests.

**A**

```
                                                     CDR H1
  CR3022: MQLVQSGTEVKKPGESLKISCKGSGYGFITYWIGWVRQMP
IGHV5-51: VQLVQSGAEVKKPGESLKISCKGSGYSFTSYWIGWVRQMP
                                          31 32 33 34 35

                      CDR H2
  CR3022: GKGLEWMGIIYPGDSETRYSPSFQGQVTISADKSINTAYL
IGHV5-51: GKGLEWMGIIYPGDSDTRYSPSFQGQVTISADKSISTAYL
          50 51 52 52a 53 54 55 56 57 58 59 60 61 62 63 64 65

                             CDR3 H3
  CR3022: QWSSLKASDTAIYYCAGGSGISTPMDVWGQGTTVTV
IGHV5-51: QWSSLKASDTAMYYCA--------------------
          95 96 97 98 99 100 100a 100b 101 102
```

**B**

```
                                         CDR L1
  CR3022: DIQLTQSPDSLAVSLGERATINCKSSQSVLYSSINKNYLA
IGKV4-1:  DIVMTQSPDSLAVSLGERATINCKSSQSVLYSSNNKNYLA
          24 25 26 27 27a 27b 27c 27d 27e 27f 28 29 30 31 32 33 34

                         CDR L2
  CR3022: WYQQKPGQPPKLLIYWASTRESGVPDRFSGSGSGTDFTLT
IGKV4-1:  WYQQKPGQPPKLLIYWASTRESGVPDRFSGSGSGTDFTLT
          50 51 52 53 54 55 56

                        CDR L3
  CR3022: ISSLQAEDVAVYYCQQYYSTPYTFGQGTKVEIK
IGKV4-1:  ISSLQAEDVAVYYCQQYYSTP------------
          89 90 91 92 93 94 95 96 97
```

**C**

```
C  A  G  G  S  G  I  S  T  P  M  D  V  W
TGTGCGGGGGGTTCGGGGATTTCTACCCCTATGGACGTCTGG
          IGHD3-10              IGHJ6

Total gene-derived nucleotides: 23
Total non-gene-derived nucleotides: 13
```

**Fig. S1. Comparison of CR3022 sequence to its putative germline sequence.** Alignment of CR3022 **(A)** with the germline IGHV5-51 sequence, and **(B)** with the germline IGKV4-1 sequence. The regions that correspond to CDR H1, H2, H3, L1, L2, and L3 are indicated. Residues that differ from the germline are highlighted in red. Residue positions in the CDRs are labeled according to the Kabat numbering scheme. **(C)** Sequence of the V-D-J junction of CR3022, with putative gene segments (blue) and N-regions (red) are indicated.

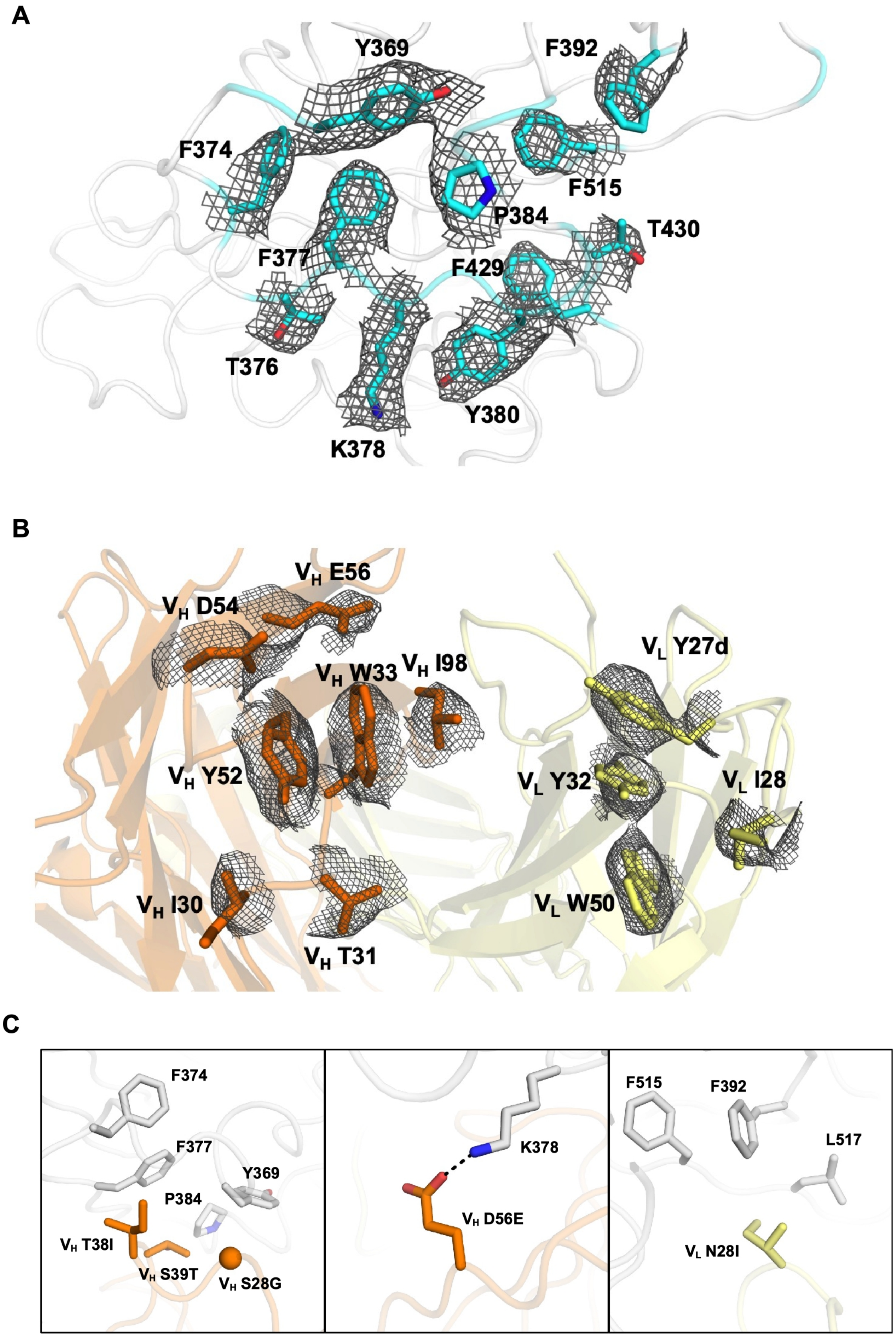


**Fig. S2. Electron density maps for epitope and paratope regions and structural**

**analysis of somatic mutations.** **(A)** Final 2Fo-Fc electron density maps for the side chains in the epitope region (cyan) of SARS-CoV-2 contoured at 1 σ. **(B)** Final 2Fo-Fc electron density maps for the paratope region of CR3022 contoured at 1 σ. The heavy chain is colored in orange, and light chain in yellow. Residues are labeled. **(C)** Somatic mutations $V_H$ S28G, $V_H$ T38I, $V_H$ S39T, $V_H$ D56E, and $V_L$ N28I are located in the CR3022 paratope region. Hydrogen bonds are represented by dashed lines. CR3022 heavy chain is in orange and light chain is in yellow. SARS-CoV-2 RBD is in light grey.

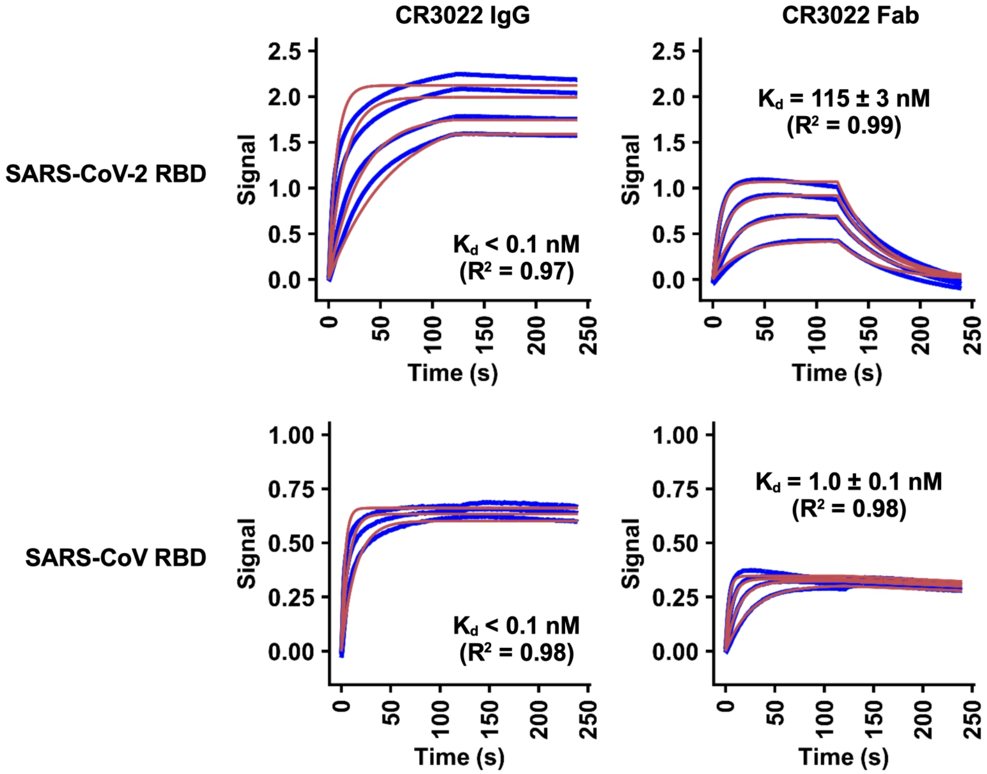


**Fig. S3. Sensorgrams for binding of CR3022 IgG and Fab to RBDs of SARS-CoV-2 and SARS-CoV.** Binding kinetics of CR3022 Fab and IgG against the RBDs of SARS-CoV-2 and SARS-CoV were measured by biolayer interferometry (BLI). Y-axis represents the response. Blue lines represent the response curves and red lines represent the 1:1 binding model. Binding kinetics were measured for three to four concentrations of IgG or Fab at 2-fold dilution ranging from 500 nM to 62.5 nM. The $K_d$ and $R^2$ of the fitting are indicated. The enhanced binding of IgG as compared to Fab is likely due to bivalent binding. Of note, stoichiometry cannot be inferred from this experiment.

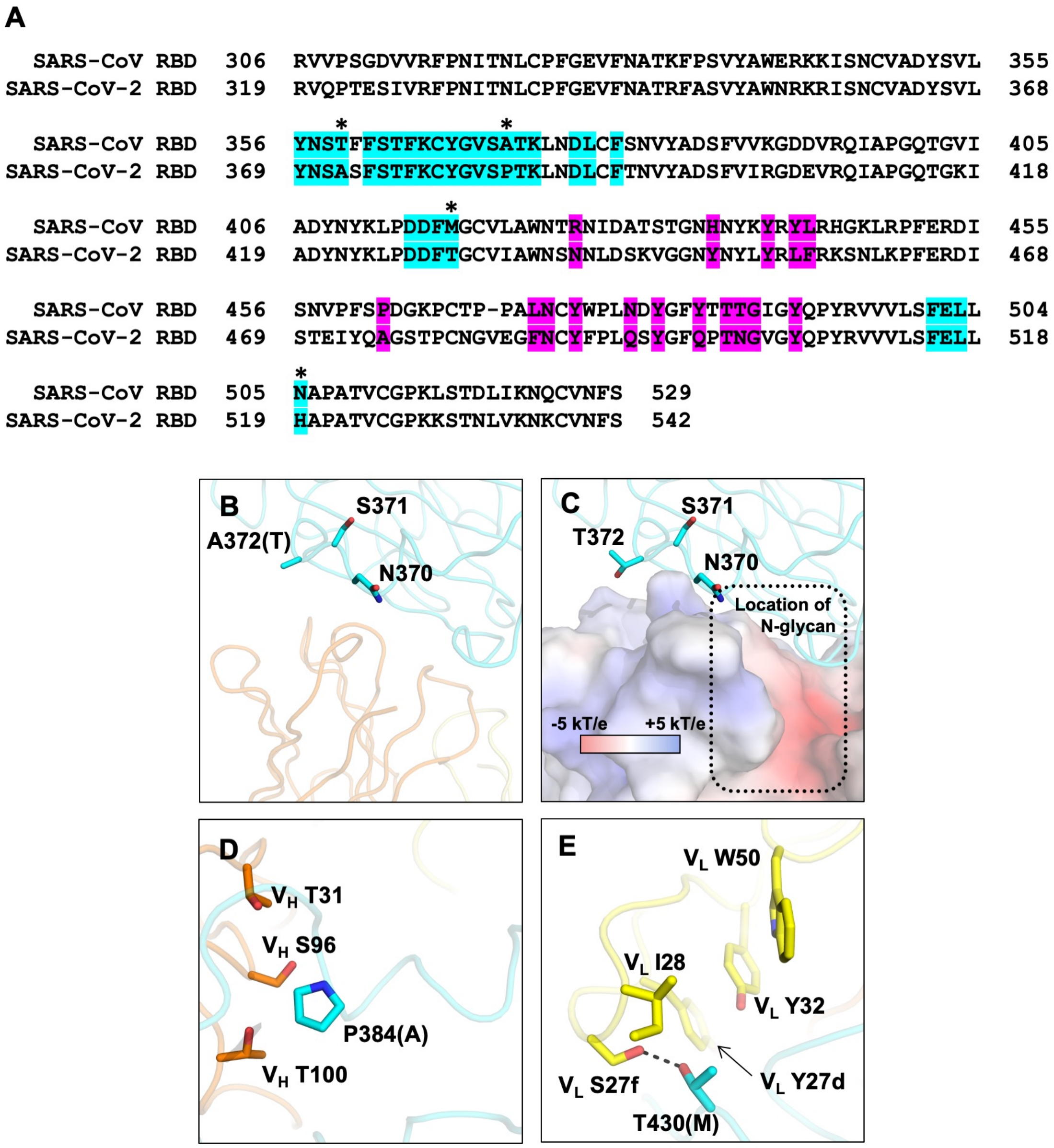


**Fig. S4. Non-conserved epitope residues. (A)** Sequence alignment of SARS-CoV-2 RBD and SARS-CoV RBD. CR3022 epitope residues are colored cyan. ACE2-binding residues are colored magenta. Non-conserved epitope residues are marked by asterisks. **(B-E)** Interactions between the non-conserved epitope residues and CR3022 are shown. Amino-acid variants observed in SARS-CoV are in parenthesis. SARS-CoV-2 RBD is colored in cyan, CR3022 heavy chain in orange, and CR3022 light chain in yellow. Residues are numbered according to their positions on the SARS-CoV-2 S protein sequence. **(B)** While SARS-CoV-2 has an Ala at residue 372, SARS-CoV has a Thr, which introduces an N-glycosylation site at residue N370. **(C)** The potential location of N370 glycan in SARS-CoV RBD is indicated by the box. CR3022 is shown as an electrostatic potential surface presentation. **(D)** P384 interacts with T31, S96, and T100 of CR3022 heavy chain. Ala at this position in SARS-CoV would allow the backbone to adopt a different conformation when binding to CR3022. **(E)** T430 forms a hydrogen bond with S27f of CR3022 light chain. Met at this position in SARS-CoV would instead likely insert its side chain into the hydrophobic pocket formed by Y27d, I28, Y32, and W50 of CR3022 light chain.

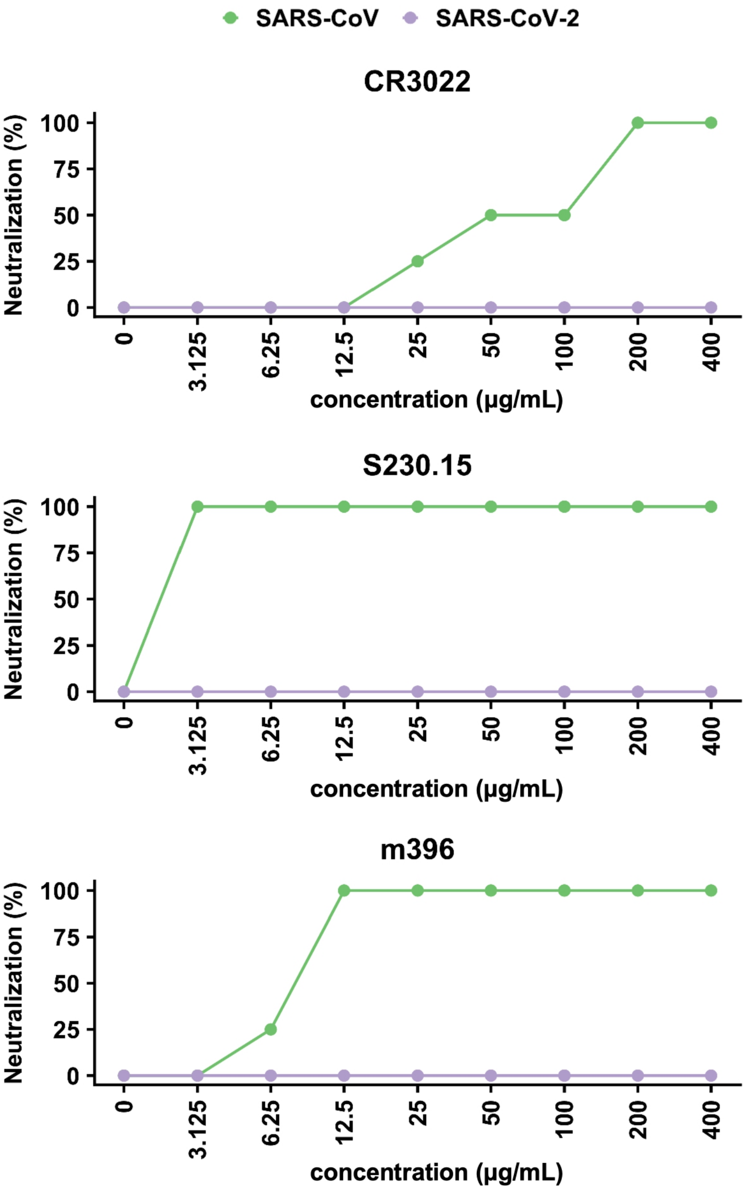


**Fig. S5. Neutralization activity of SARS-neutralizing monoclonal antibodies against SARS-CoV-2 and SARS-CoV.** Microneutralization of SARS-CoV-2 and SARS-CoV with monoclonal antibodies **(A)** CR3022, **(B)** S230.15 (*39*), and **(C)** m396 (*16*). Neutralization of 100 $TCID_{50}$ of each virus was performed in quadruplicate.

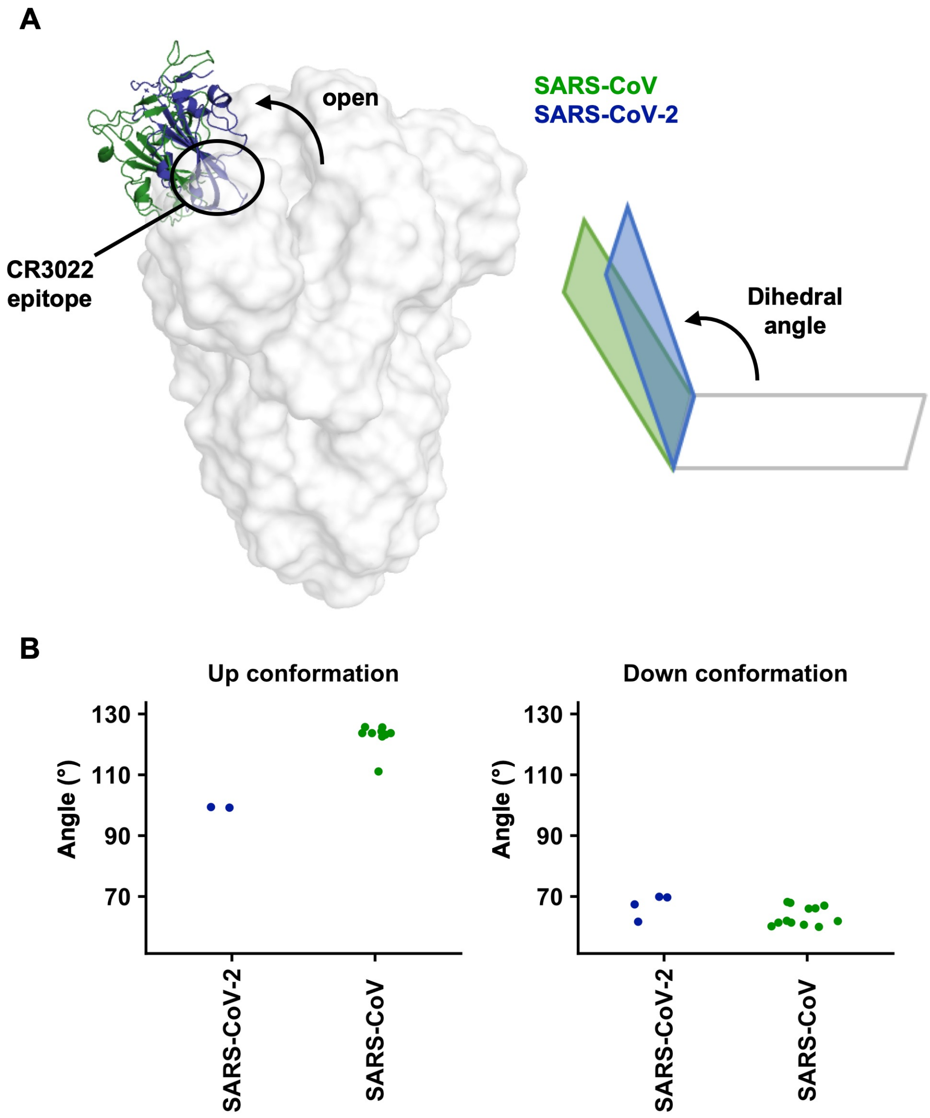


C

| PDB | 2AJF (copy 1) | 2AJF (copy 2) | 6VSB (protomer 1) | 6VSB (protomer 2) | 6VSB (protomer 3) | 6VYB (protomer 1) | 6VYB (protomer 2) | 6VYB (protomer 3) |
|---|---|---|---|---|---|---|---|---|
| State | - | - | up | down | down | up | down | down |
| RMSD (Å) | 1.22 | 1.54 | 1.11 | 1.94 | 1.00 | 1.07 | 1.37 | 1.11 |

**Fig. S6. Comparison of the "up" conformations between the SARS-CoV-2 and SARS-CoV S proteins. (A)** Structural alignment of the one-"up" S proteins from SARS-CoV-2 (PDB 6VSB (*17*)) and SARS-CoV (PDB 6CRW (*21*)). The RBD with an "up" conformation from SARS-CoV-2 is in blue and from SARS-CoV is in green. **(B)** The RBD open angle is represented by a dihedral angle between the RBD and the horizontal plane. Given that the RBD of interest is on protomer 1, the dihedral angle is measured by the angle of P507 Cα (protomer 1) – R983 Cα (protomer 2) – R983 Cα (protomer 1) – R983 Cα (protomer 3) for SARS-CoV-2, which corresponds to of P493 Cα (protomer 1) – R965 Cα (protomer 2) – R965 Cα (protomer 1) – R965 Cα (protomer 3) for SARS-CoV. For this analysis, S protein structures with PDB IDs of 6VSB (*17*), 6VYB (*18*) were used for SARS-CoV-2, whereas 5X5B (*19*), 6CRX (*21*), 6VS0, 6VS1, 6CRW (*21*), 6ACD (*47*), and 6CRZ (*21*) were used for SARS-CoV. **(C)** Root-mean-square deviation (RMSD Cα) of the RBDs from different structures were computed using "super" function in PyMOL with no refinement cycle. PDB 2AJF (*42*), 6VSB (*17*), and 6VYB (*18*) were used in this analysis.

### Table S1. X-ray data collection and refinement statistics

| | |
|---|---|
| **Data collection** | |
| Beamline | APS 23ID-D |
| Wavelength (Å) | 1.03322 |
| Space group | $P4_122$ |
| Unit cell parameters | a=b=147.5°, c=200.2° |
| Resolution (Å) | 50.0-3.10 (3.21-3.10)[a] |
| Unique reflections | 41,206 (938)[a] |
| Redundancy | 6.7 (5.5)[a] |
| Completeness (%) | 100.0 (100.0)[a] |
| $<I/\sigma_I>$ | 18.8 (1.0)[a] |
| $R_{sym}$[b] (%) | 13.6 (>100)[a] |
| $R_{pim}$[b] (%) | 4.0 (54.0)[a] |
| $CC_{1/2}$[c] (%) | 100.0 (59.1)[a] |
| **Refinement statistics** | |
| Resolution (Å) | 50.0-3.10 |
| Reflections (work) | 41,137 |
| Reflections (test) | 2,030 |
| $R_{cryst}$[d] / $R_{free}$[e] (%) | 22.3/24.3 |
| No. of atoms | 4,936 |
| Macromolecules | 4,906 |
| Ligands | 30 |
| Average *B*-value ($Å^2$) | 99 |
| Macromolecules | 99 |
| Ligands | 137 |
| Wilson *B*-value ($Å^2$) | 95 |
| **RMSD from ideal geometry** | |
| Bond length (Å) | 0.002 |
| Bond angle ($^o$) | 0.52 |
| **Ramachandran statistics (%)** | |
| Favored | 97.2 |
| Outliers | 0.0 |
| **PDB code** | 6W41 |

[a] Numbers in parentheses refer to the highest resolution shell.

[b] $R_{sym} = \Sigma_{hkl} \Sigma_i | I_{hkl,i} - <I_{hkl}> | / \Sigma_{hkl} \Sigma_i I_{hkl,i}$ and $R_{pim} = \Sigma_{hkl} (1/(n-1))^{1/2} \Sigma_i | I_{hkl,i} - <I_{hkl}> | / \Sigma_{hkl} \Sigma_i I_{hkl,i}$, where $I_{hkl,i}$ is the scaled intensity of the i[th] measurement of reflection h, k, l, $<I_{hkl}>$ is the average intensity for that reflection, and *n* is the redundancy.

[c] $CC_{1/2}$ = Pearson correlation coefficient between two random half datasets.

[d] $R_{cryst} = \Sigma_{hkl} | F_o - F_c | / \Sigma_{hkl} | F_o | \times 100$, where $F_o$ and $F_c$ are the observed and calculated structure factors, respectively.

[e] $R_{free}$ was calculated as for $R_{cryst}$, but on a test set comprising 5% of the data excluded from refinement.

**Table S2. Hydrogen bonds and salt bridges identified at the SARS-CoV-2 RBD and CR3022 interface using the PISA program**

| **Hydrogen bonds** | | |
|---|---|---|
| **SARS-CoV-2 RBD** | **Dist. (Å)** | **CR3022** |
| PHE377[N] | 3.0 | $V_H$ TYR52[OH] |
| LYS378[N] | 3.9 | $V_H$ TYR52[OH] |
| LYS378[NZ] | 2.7 | $V_H$ ASP54[OD2] |
| LYS378[NZ] | 2.4 | $V_H$ GLU56[OE1] |
| LYS386[NZ] | 3.3 | $V_H$ ASP101[OD1] |
| PHE377[O] | 2.5 | $V_H$ TYR52[OH] |
| CYS379[O] | 3.2 | $V_H$ ILE98[N] |
| THR430[OG1] | 2.4 | $V_L$ SER27f[OG] |
| GLY381[O] | 2.3 | $V_L$ TYR32[OH] |
| **Salt bridges** | | |
| LYS378[NZ] | 2.7 | $V_H$ ASP54[OD2] |
| LYS378[NZ] | 2.4 | $V_H$ GLU56[OE1] |
| LYS378[NZ] | 3.4 | $V_H$ ASP54[OD1] |
| LYS386[NZ] | 3.3 | $V_H$ ASP101[OD1] |
| LYS386[NZ] | 3.6 | $V_H$ ASP101[OD2] |

**Table S3. Epitope residues on the SARS-CoV-2 RBD and their buried surface area upon binding to CR3022**

| SARS-CoV-2 RBD | BSA (Å$^2$) |
|---|---|
| TYR369 | 50 |
| ASN370 | 41 |
| SER371 | 5 |
| ALA372 | 11 |
| PHE374 | 17 |
| SER375 | 17 |
| THR376 | 23 |
| PHE377 | 63 |
| LYS378 | 96 |
| CYS379 | 35 |
| TYR380 | 66 |
| GLY381 | 11 |
| VAL382 | 8 |
| SER383 | 37 |
| PRO384 | 33 |
| THR385 | 42 |
| LYS386 | 105 |
| ASP389 | 6 |
| LEU390 | 32 |
| PHE392 | 17 |
| ASP427 | 7 |
| ASP428 | 58 |
| PHE429 | 7 |
| THR430 | 50 |
| PHE515 | 6 |
| GLU516 | 6 |
| LEU517 | 49 |
| HIS519 | 20 |

### REFERENCES


38. D. C. Ekiert *et al.*, A highly conserved neutralizing epitope on group 2 influenza A viruses. *Science* **333**, 843-850 (2011).

39. A. C. Walls *et al.*, Unexpected receptor functional mimicry elucidates activation of coronavirus fusion. *Cell* **176**, 1026-1039 e1015 (2019).

40. Z. Otwinowski, W. Minor, Processing of X-ray diffraction data collected in oscillation mode. *Methods Enzymol* **276**, 307-326 (1997).

41. A. Teplyakov *et al.*, Antibody modeling assessment II. Structures and models. *Proteins* **82**, 1563-1582 (2014).

42. F. Li, W. Li, M. Farzan, S. C. Harrison, Structure of SARS coronavirus spike receptor-binding domain complexed with receptor. *Science* **309**, 1864-1868 (2005).

43. K. Arnold, L. Bordoli, J. Kopp, T. Schwede, The SWISS-MODEL workspace: a web-based environment for protein structure homology modelling. *Bioinformatics* **22**, 195-201 (2006).

44. P. Emsley, K. Cowtan, Coot: model-building tools for molecular graphics. *Acta Crystallogr D Biol Crystallogr* **60**, 2126-2132 (2004).

45. P. D. Adams *et al.*, PHENIX: a comprehensive Python-based system for macromolecular structure solution. *Acta Crystallogr D Biol Crystallogr* **66**, 213-221 (2010).

46. N. C. Wu *et al.*, In vitro evolution of an influenza broadly neutralizing antibody is modulated by hemagglutinin receptor specificity. *Nat Commun* **8**, 15371 (2017).

47. W. Song, M. Gui, X. Wang, Y. Xiang, Cryo-EM structure of the SARS coronavirus spike glycoprotein in complex with its host cell receptor ACE2. *PLoS Pathog* **14**, e1007236 (2018).
